# Past human expansions shaped the spatial pattern of Neanderthal ancestry

**DOI:** 10.1101/2022.12.22.521596

**Authors:** Claudio S. Quilodrán, Jérémy Rio, Alexandras Tsoupas, Mathias Currat

**Affiliations:** Department of Genetics and Evolution, University of Geneva, Switzerland; Department of Biology, University of Fribourg, Switzerland; Institute of Genetics and Genomics in Geneva (IGE3), University of Geneva, Switzerland

**Author notes:** contributed equally to this work (Claudio S. Quilodrán, Jérémy Rio).

**Keywords:** Population expansion, Admixture, Paleogenomics, Evolutionary dynamics, Introgression, Neanderthal ancestry

## Abstract

The worldwide expansion of modern humans (*Homo sapiens*) from Africa started before the extinction of Neanderthals (*Homo neanderthalensis*). Both species coexisted and interbred, as revealed by the sequencing of Neanderthal genomes, leading to ~2% Neanderthal DNA in modern Eurasians^1,2^, with slightly higher introgression in East Asians than in Europeans^3–6^. These distinct levels of ancestry have been argued to result from selection processes^7,8^. However, recent theoretical simulations have shown that range expansions could be another explanation^9,10^. This hypothesis would lead to the generation of spatial gradients of introgression, increasing with the distance from the source of the expansion, i.e., Africa for modern humans. Here, we investigate the presence of Neanderthal introgression gradients after past human expansions by analysing an extended palaeogenomic dataset of Eurasian populations. Our results show that the Out-of-Africa expansion of modern humans into Eurasia resulted in spatial gradients of Neanderthal ancestry that persisted through time. Moreover, while keeping the same gradient orientation, the expansion of early Neolithic farmers into western Eurasia contributed decisively to reducing the average level of Neandertal genomic introgression in European compared to Asian populations. This is because Neolithic farmers carried less Neanderthal DNA than preceding Palaeolithic hunter-gatherers. This study shows that inferences about past population dynamics within our species can be made from the spatiotemporal variation in archaic introgression.

## Introduction

Sequencing of Neanderthal genomes has revealed that ~2% of the DNA of modern humans (MHs) outside of Africa is more similar to DNA from Neanderthals (NEs) than it is to DNA from sub-Saharan populations^1,2^. Two main, nonexclusive hypotheses have been proposed to explain this pattern: (i) hybridization between NEs and MHs during their expansion out of Africa (OOA), leading to the introgression of Neanderthal DNA segments into modern humans^1,2^; and (ii) incomplete lineage sorting resulting from ancestral population structure in Africa, with ancestors of non-Africans more closely related to NEs than to sub-Saharan Africans^11^. Evidence in favour of hybridization has accumulated during the last decade^5,12,13^. However, the number, timing, and locations of interbreeding events between MH and NE remain unclear. While early studies have suggested a single hybridization pulse in the Middle East^1,2^, a growing body of research supports the hypothesis of multiple hybridization events^6,10,14–16^. In particular, it has been shown that multiple hybridisation events over time and space in western Eurasia are consistent with the levels of Neanderthal ancestry observed in modern populations.^10^.

While NE ancestry is relatively uniform among modern Eurasian populations^1,2^, it is approximately 8-24% higher in East Asia than in Europe^3–6^. This observation is surprising since the known distribution of NEs was almost exclusively in the western part of Eurasia^17^. Three major hypotheses have been proposed to explain the difference in NE ancestry between western and eastern Eurasian populations: (i) higher effective population size in Europeans compared to Asians, which led to a stronger effect of purifying selection acting on deleterious NE alleles in the former^7^; (ii) dilution of NE ancestry in Europeans due to an input from a hypothetical basal or “ghost” population with little or no NE ancestry^18,19^; and (iii) multiple pulses of NE introgression, where the original Eurasian introgression pulse was supplemented by additional pulses after the divergence between the European and Asian populations, resulting in different NE ancestry levels^6,8,15,19^.

Recently, an additional hypothesis has been proposed, where different levels of NE ancestry between western Europe and Eastern Asia are the result of the range expansion of MHs after the OOA event^9^. Population range expansions have important evolutionary consequences, including (i) creating gradients of allele frequencies^20,21^; (ii) increasing the frequency of specific alleles, whether neutral^22,23^ or under natural selection^24,25^; (iii) decreasing genetic diversity along the axis of expansion^26,27^; and (iv) increasing mutational load in populations through the maintenance of deleterious alleles^28^. In addition, when admixture with a local population occurs, population expansions tend to disproportionately increase the genetic contribution of the local population to the invasive genetic pool, even if interbreeding is limited^29^. This latter effect is expected to result in the formation of spatial gradients of neutral introgression along the axis of the biological invasions with hybridization (see Fig. 1A). Under this assumption, introgression of local genes (i.e., NEs) increases in the invasive population (i.e., MHs) with the distance from the source of the expansion (i.e., Africa). This is due to the combined effects of i) demographic imbalance between the expanding population and the local population at demographic equilibrium, ii) continuous hybridization events at the wave front of the range expansion, and iii) genetic surfing. This hypothesis could therefore explain different levels of Neanderthal ancestry in Europe and East Asia by differences in geographical distance from the source of the MH expansion in Africa^9^. Quilodrán, et al.^9^ showed that the process of range expansion could explain the difference in NE ancestry between Europe and East Asia using computational simulations based on a limited amount of genomic information from the two extreme sides of the continent (West and East). However, neither a detailed inspection of geographic introgression patterns (i.e., the existence of gradients) nor their change over time was included in this preliminary study. Indeed, range expansions occurred not only during the OOA expansion^10^ but also during other prehistoric periods^30^. This includes the European Neolithic transition, when farmers coming from southeast Europe partially replaced hunter-gatherers^31–34^, as well as the Bronze Age, with the spread of pastoralist populations from Eurasian steppes^35–37^. Therefore, multiple population movements during recent human history could have contributed to shaping NE ancestry across time and space because distinct expanding populations can carry various levels of NE ancestry^20,30,38^.

**Figure 1:**
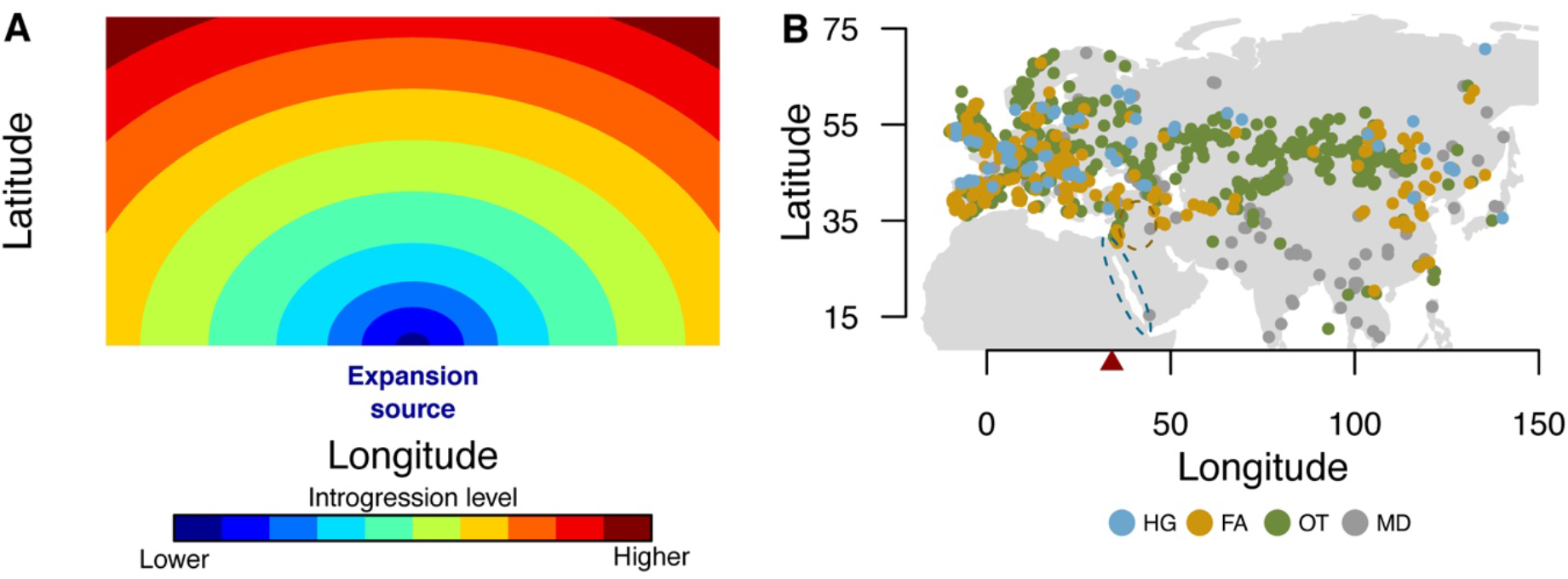
A) Schematic representation of the expected spatial gradient of introgression from the local taxon into an invasive taxon after a biological invasion with hybridization. B) Distribution of samples in Eurasia used for elucidating spatial gradients of Neanderthal ancestry in modern humans. The coloured dots represent palaeogenomic samples of hunter-gatherers (HG, ~40000–6000 years BP, *n* = 129), early farmers (FA, ~10000–2000 years BP, *n* = 679), other ancient (OT, ~6400–300 years BP, *n* = 1726), and modern (MD, current time, *n* = 91). The dotted ellipse represents the presumed geographic source of the Palaeolithic out of Africa expansion into Eurasia (~50000 years BP), and the dotted circle represents the source of the Neolithic expansion of agricultural populations from the fertile crescent (~11000 years BP). The red triangle represents the longitudinal limit (34°) that, in our study, separates European (*n*=1517) from Asian (*n*=1108) population samples.

Here, we investigate whether spatial gradients of introgression consistent with the range expansion hypothesis have occurred in Eurasia by examining the levels of NE ancestry in human populations distributed across space and time. Our study demonstrates that spatiotemporal levels of introgression provide valuable information about past population dynamics, suggesting multiple episodes of range expansion as a main force shaping archaic introgression levels during human evolutionary history.

## Results and discussion

### Spatial gradients of Neanderthal ancestry in Eurasia

We analysed an extended dataset of 4464 published ancient and modern genomes (from ~40000 years BP to present time) retrieved from the Allen Ancient DNA database^39^. We associated each genome with one of the following population groups: Palaeolithic/Mesolithic hunter-gatherers (HGs), Neolithic Chalcolithic farmers (FAs), other ancient samples (OTs) or modern samples (MDs). We estimated NE introgression for all genomes and averaged the introgression estimates for genomes from the same geographic coordinates, time periods and population group, leading to *n* = 2625 samples (Fig. 1B). We then explored the fixed effect of latitude, longitude, time (dates in years BP), continental area (Europe or Asia), and their interactions by using a linear mixed model (LMM) with log-transformed NE ancestry as the response variable. LMMs are particularly useful for dealing with hierarchical structures and the nonindependence of the dataset. We investigated the random effect of the population group, the period nested within these groups (in clusters of 500 years), and the spatial and temporal autocorrelation of data. This model was called “Full Eurasia” because it uses the whole palaeogenomic dataset. Based on the lowest AIC value^40^, the best LMM was retained (Table S1, supporting information).

By considering the average date of all samples as a time reference (~4200 years BP), we observed a linear relationship of NE ancestry with latitude and longitude, both in Europe and Asia (Fig. 2, Table S1). These geographic patterns support the hypothesis of spatial introgression gradients generated after population expansion with hybridization^10^, schematically represented in Fig. 1A, in which the introgression of local genes (i.e., NEs) is expected to increase in the invasive population (i.e., MHs) with the distance from the source of its expansion (i.e., Africa). While a positive relationship with latitude is observed in Europe and Asia (Fig. 2A), a contrasting relationship is observed with longitude, positive in Asia and negative in Europe (Fig. 2B). The increasing NE introgression with latitude is compatible with the OOA expansion of MHs from southern to northern areas of Eurasia while hybridizing with NEs. The longitudinal pattern is compatible with a source of expansion in the Middle East, from which NE ancestry is expected to increase with longitude in Asia but decrease with longitude in Europe. Although alternative evolutionary forces may also be responsible for creating introgression gradients (e.g., spatially varying selective pressure), the specific pattern we observed with a source of all spatial gradients in the Middle East makes population range expansion the most parsimonious explanation.

**Figure 2.**
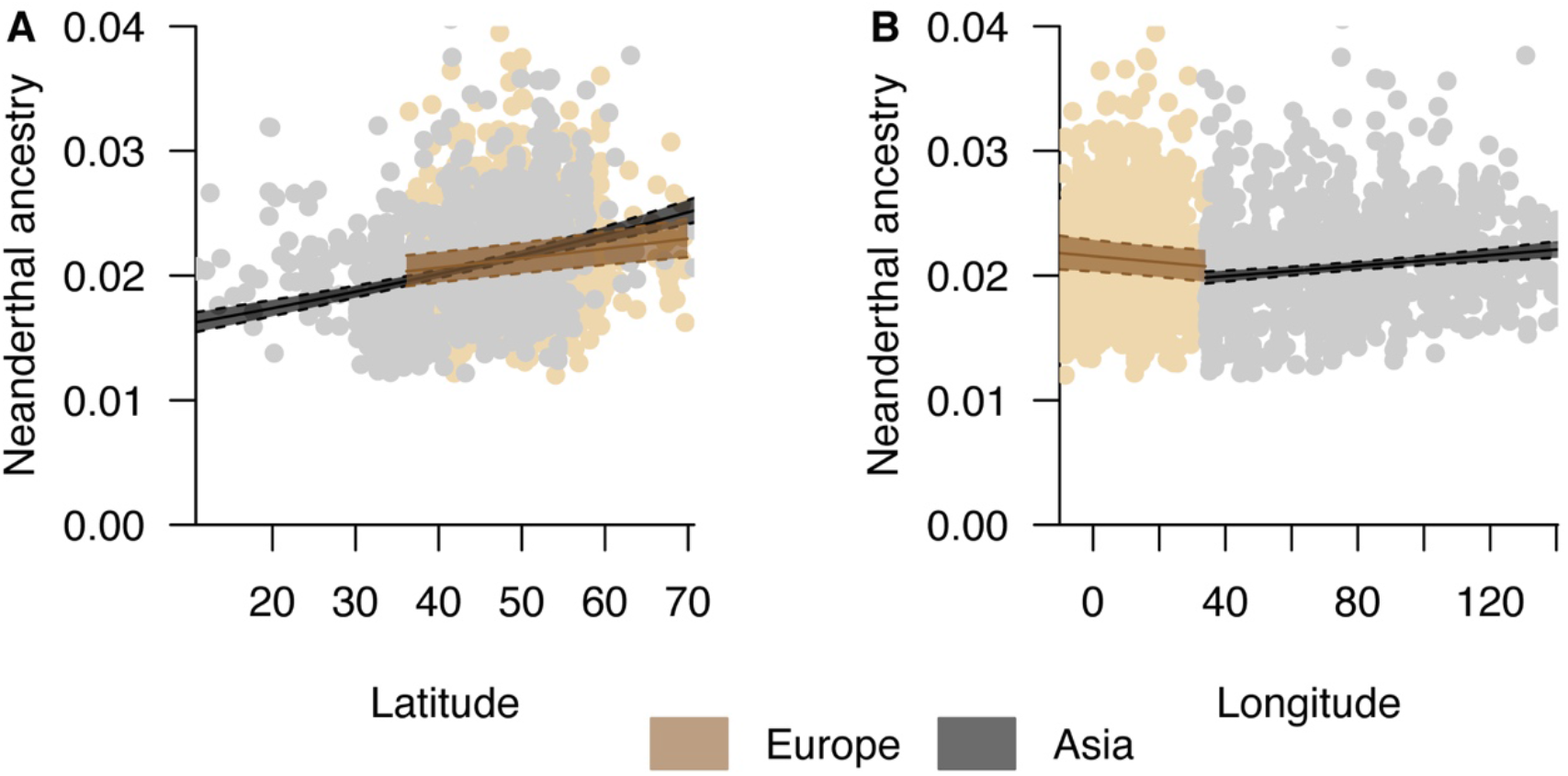
Effects of latitude and longitude on the level of Neanderthal ancestry in both Europe and Asia. The solid and dotted lines represent the estimated values and 95% confidence intervals, respectively. The coloured dots represent the distribution of the full dataset of ancient and modern DNA samples used in the “Full Eurasia” analysis (*n* = 2625).

Alternatively, spatial variation of NE ancestry in MH could result from an unequal distribution of NEs in Eurasia, with more interspecific interactions occurring in areas where NE were more numerous, resulting in more local admixture. We find a higher NE ancestry level in Europe than in Asia after the OOA (Fig. 3), which is in accordance with the current fossil record of NEs in Eurasia, with more accumulated evidence in Europe. Moreover, our results showing more NE ancestry in northern Eurasia than in southern Eurasia further concur with evidence of NE presence in the northern Himalayas^17^, even if an undetected presence in the south cannot be ruled out. Nevertheless, even in the case of unequal distribution of the local species, increasing gradients of introgression in the invasive population (i.e., MHs) resulting from range expansion may remain a valid explanation. For instance, this is expected after a range expansion where the local population is only occupying a part of the area^9^, as was probably the case for NEs in Eurasia.

**Figure 3.**
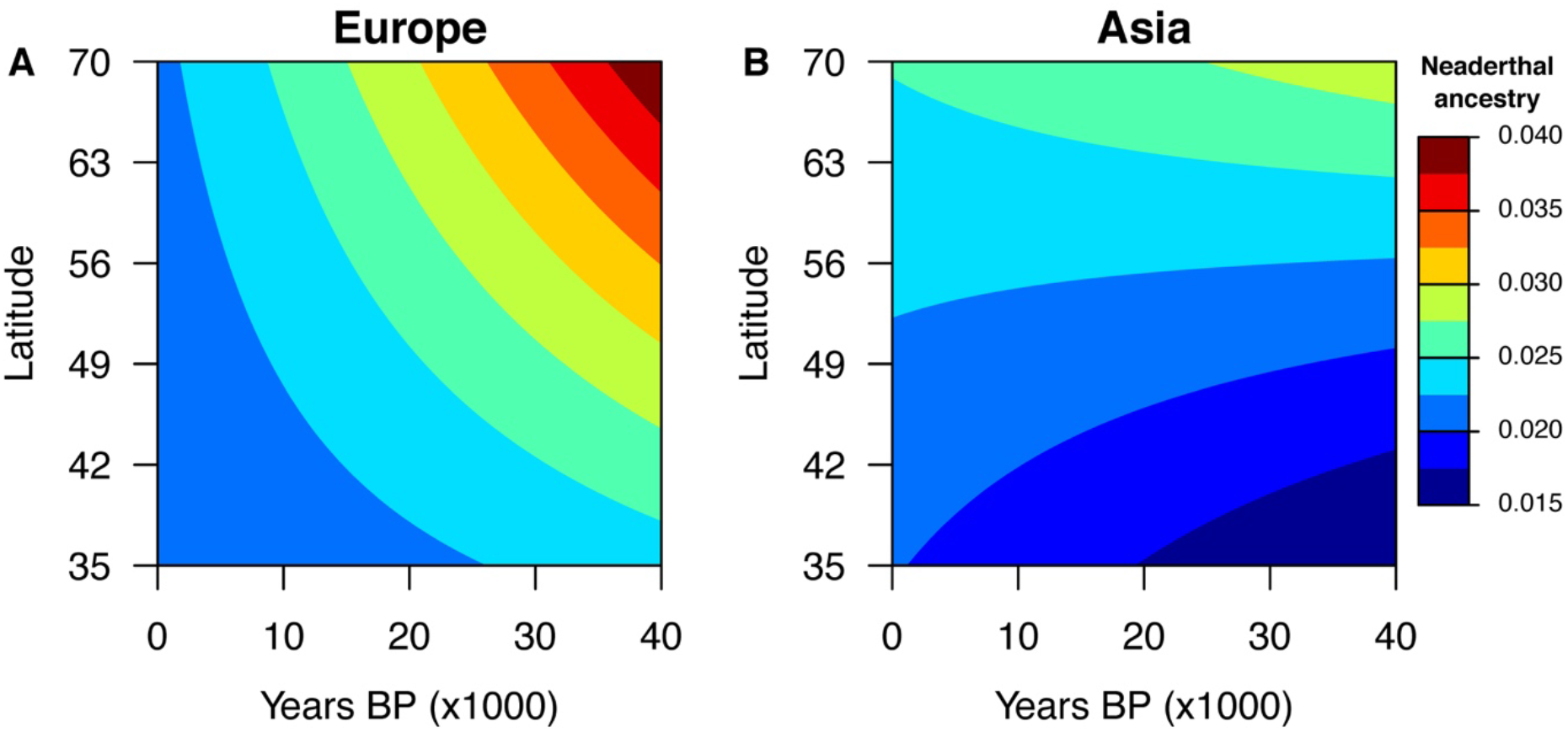
Influence of latitude and time on the level of Neandertal ancestry in both Europe and Asia. The analysis considers the full dataset of ancient and modern DNA samples (“Full Eurasia”, *n* = 2625). The y-axis corresponds to the range where both regions (Asia and Europe) have the most data.

However, our analysis does not support the hypothesis that a slightly higher level of NE ancestry in East Asia than in Western Europe today could be explained by the greater distance to the source of the MH expansion in Asia than in Europe^9^. This hypothesis was made from modern DNA data only, while our results also considered ancient DNA samples. Using palaeogenomic data, we show the reverse pattern for samples older than 20000 years BP, with more NE ancestry observed in Europe than in Asia (Fig. 3). The current pattern of NE ancestry being higher in Asia than in Europe must thus have developed at a later stage.

### Temporal variation in Neanderthal ancestry

Our results suggest that the longitudinal gradient slope has remained similar over the last 40000 years (Table S1, supporting information), whereas the latitudinal gradient of NE ancestry has significantly changed over time (F = 4.4, P = 0.03). The latitudinal pattern is more prominent during the early period in Europe and becomes less visible ~30000 years BP, together with an overall reduction in NE ancestry (Fig. 3). While this implies that the level of NE ancestry in Eurasia may not have been uniformly distributed across space as is observed today, this expectation needs to be confirmed with more palaeogenomes because the interaction between latitude and time is no longer significant when considering the average of all candidate models (based on a cumulative weighted AIC of 90%, *ωw_i_* ≥ 0.90, Table S2, supporting information) instead of the best model only.

The variation in NE ancestry across time is currently debated. It has been proposed that ancient European genomes showed more NE ancestry compared to present-day Europeans^12^, but this result was questioned because of a methodological bias in the ancestry estimation procedure^41^. Nevertheless, it has been estimated that NE ancestry could have been as high as 10% at the time of admixture before decreasing rapidly to the current rate of ~2%^42^. Here, we show that the temporal reduction in NE ancestry is linked to latitude. The southern samples in Europe show an almost constant NE ancestry through time, while the northern samples experienced a reduction between approximately 40000 and 20000 years BP. The latitudinal gradient could have undergone important changes, possibly due to population expansions and contractions experienced by MHs during the Last Glacial Maximum (LGM) or other, more limited ice ages. Our results show that this gradient becomes less evident in modern data (Fig. 3). Because the longitudinal pattern has been less affected during the last 40000 years, it may represent a relict signature of the OOA range expansion that occurred during MH evolution between 60000 and 45000 years BP.

Natural selection has been invoked to explain the reduction in NE ancestry over time^7,43^, but different historical processes could have also played an important role. This includes population expansions and contractions^44,45^, as well as interactions between genetically differentiated populations with different levels of NE ancestry^18,19^. In Europe, a prominent genetic transition occurred during the spread of early Neolithic farmers, when they partially replaced Palaeolithic/Mesolithic hunter-gatherers (e.g. the so-called Neolithic transition)^31,32–34^. This transition began in the Fertile Crescent ~11000 years BP^46^, and its consequences on the distribution of NE ancestry have been weakly explored thus far^18^.

### Early farmers carried less Neanderthal ancestry than hunter-gatherers in Europe

We thus explored more specifically the variation in the level of NE ancestry across time and population groups (HGs, FAs, OTs, *n* = 2534 in total). The OT group includes all ancient samples that are neither FAs nor HGs, including for example, the Bronze Age and more recent periods. Samples from the MD group were excluded from this analysis because they do not allow us to explore temporal variation in NE ancestry (all modern data are associated with the same date). We included population groups, continental area (either Europe or Asia), time and their interactions as fixed variables, also correcting for spatial autocorrelation in the dataset. This model is called “Ancient Eurasia” because it only considers ancient samples and its best LMM is presented in Table S1 (supporting information). We observed an influence of time on the differences between Europe and Asia (F = 9.22, P < 0.01) and population groups (F = 9.41, P < 0.01), with an overall higher NE ancestry level for HGs than for FAs, particularly visible in Europe (Fig. 4). At approximately 10000 years BP, when the first FA appeared in the Near East, the difference between FAs and HGs was significant in Europe (HG: 0.024 ± 0.001 (estimated mean ± se), FA: 0.019 ± 0.001, t ratio = −6.14, P < 0.001), as well as in Asia (HG: 0.022 ± 0.001, FA: 0.018 ± 0.001, t ratio = −6.14, P < 0.001). Approximately 6000 years BP, when farming was well established but some HG populations persisted, the difference in NE ancestry was still significant between the HG (0.023 ± 0.001) and FA (0.020 ± 0.0002) populations in Europe (t ratio = −4.21, P < 0.001), as well as between the HG and OT (0.020 ± 0.0003) populations (t ratio = 3.51, P = 0.001), but not between the FA and OT populations (t ratio = −0.41, P = 0.91). A similar situation was observed in Asia at this time (FAs vs. HGs, t ratio = −4.26, P < 0.001; HGs vs. OTs, t ratio = 3.51, P = 0.001; FAs vs. OTs, t ratio = −0.41, P = 0.91). Overall, this means that earlier FA carried less NE ancestry than the former HGs of the same area. This difference vanished over time, since the level of NE ancestry in FAs increased during the cohabitation period with HGs in both geographic regions (Fig. 4). While Asian FAs reached an average level similar to that of HGs, European FAs did not reach such a high level (Fig. 4). Thus, FAs could have acted as a population that diluted NE ancestry in Western Eurasia, as previously suggested^18,19^.

**Figure 4.**
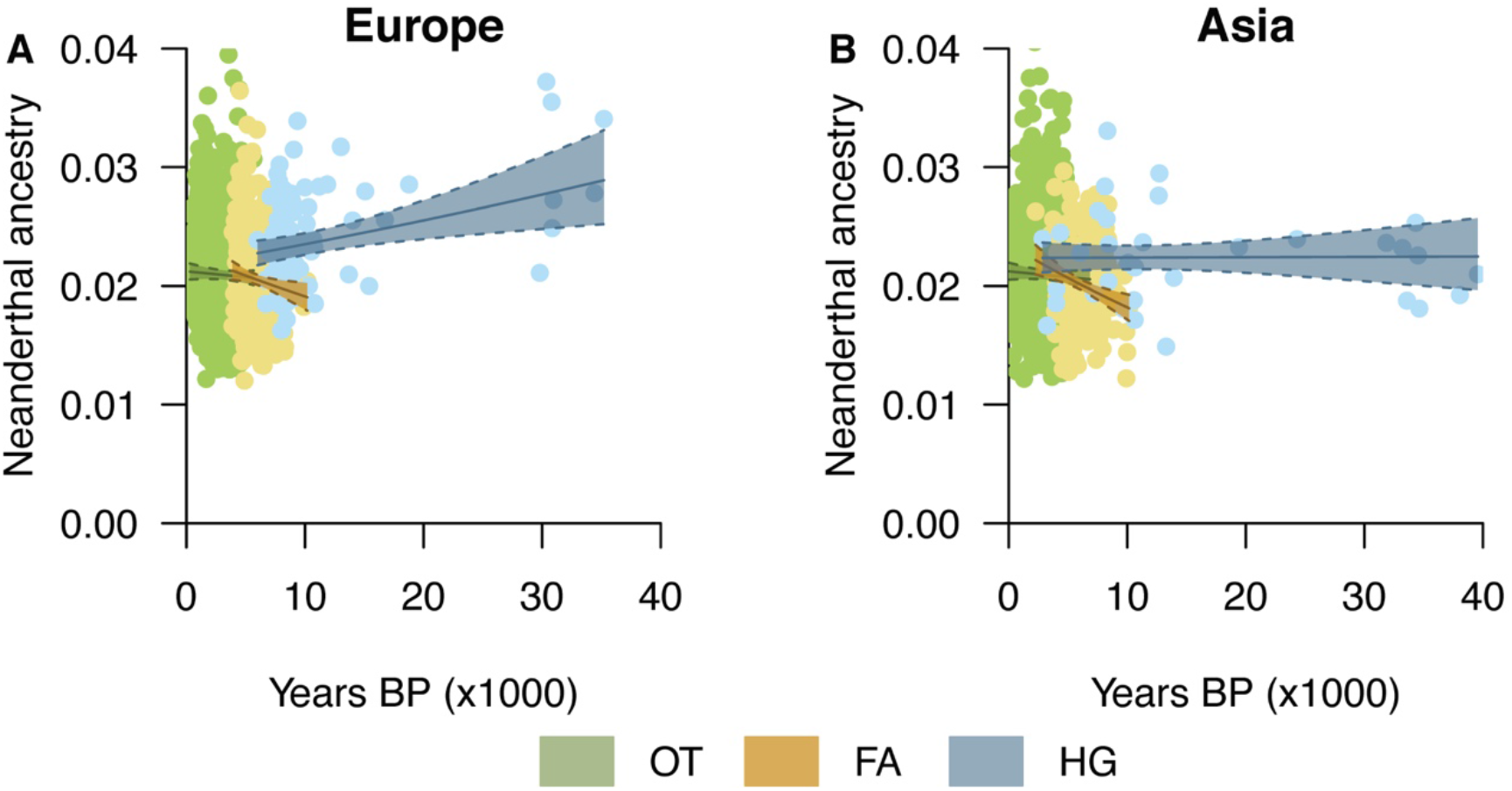
Temporal variation in the level of Neanderthal ancestry in different cultural populations of Eurasia. HGs: hunter gatherers, FAs: Neolithic farmers, OTs: other ancient samples. The solid and dotted lines represent the estimated values and 95% confidence intervals, respectively. The coloured dots represent the distribution of ancient DNA samples used in the best “Ancient Eurasia” analysis (*n* = 2534).

Multiple episodes of range expansion of populations where levels of NE ancestry differed could provide an explanation for the spatiotemporal change in NE ancestry. Our results support our former expectation that human past range expansions (i.e., HGs then FAs) contributed to the creation of spatial gradients of NE ancestry, with the level increasing from their source in Southwest Asia (Fig. 2 and Fig. 3). The second range expansion into Western Eurasia, that of early FAs with a lower level of NE ancestry, is critical to explain the overall dilution of NE ancestry in this area. The later expansion of the steppe pastoralists does not seem to have had as much influence as there is not a significant difference between the FA and OT population groups, but this would require a more detailed examination, as our OT group includes populations from different cultural periods.

### Spatial gradients in European farmers and hunter-gatherers

We then decided to focus our analysis on Europe due to the larger density and more uniform distribution of palaeogenomes than in Asia, where the Palaeolithic and Neolithic eras are represented by a lower number of samples that cover a larger area (1517 samples in Europe vs. 1108 samples in Asia for an area four times as large, Fig. 1B). Moreover, domesticated plants and animals occured in more than one site in Asia^47,48^, making exploration of past FA population dynamics more challenging than in Europe.

By using a subsample of data restricted to Europe (*n* = 1517), we explored the effect of latitude, longitude, and population groups (HG-FA-OT-MD) on NE ancestry, controlling for the fixed effect of time. We excluded cross-level interactions between time and population groups because of the lack of temporal variation in MD samples. The LMM is called “Europe” and its best version is presented in Table S1 (supporting information). The interaction between longitude and population group was nonsignificant and excluded during model selection, together with the interaction between latitude and population groups (Table S1 and Table S2). This implies that the negative longitudinal trend and positive latitudinal trend shown in Figure 2 remain for all population groups in Europe, despite the higher NE ancestry in HGs compared to other cultural populations (Fig. 5). The interaction between latitude and longitude was significant (F = 29.29, P < 0.001), meaning that their introgression slopes were interdependent for all population groups (Fig. 5 and Fig. S1, supporting information). When considering the full model that includes all fixed variable interactions, the only spatial gradient that has changed is the slope of latitude for HGs compared to FAs (F = 2.68, P = 0.04, Table S3, supporting information). Although the result for HGs may be influenced by the scarcity of samples during the Palaeolithic, this analysis suggests that the latitudinal variation could have changed more than the longitudinal variation during this period (Fig. 3 and Fig. 5). Different events of population contractions and expansions during the Palaeolithic related to the LGM^44,45^ could have affected the latitudinal gradient more than the one seen for longitude. Overall, the spatial trend remains similar across different periods of time, becoming less pronounced in modern samples (Fig. S1, supporting information) and with a higher NE ancestry in HGs (Fig. 5).

**Figure 5.**
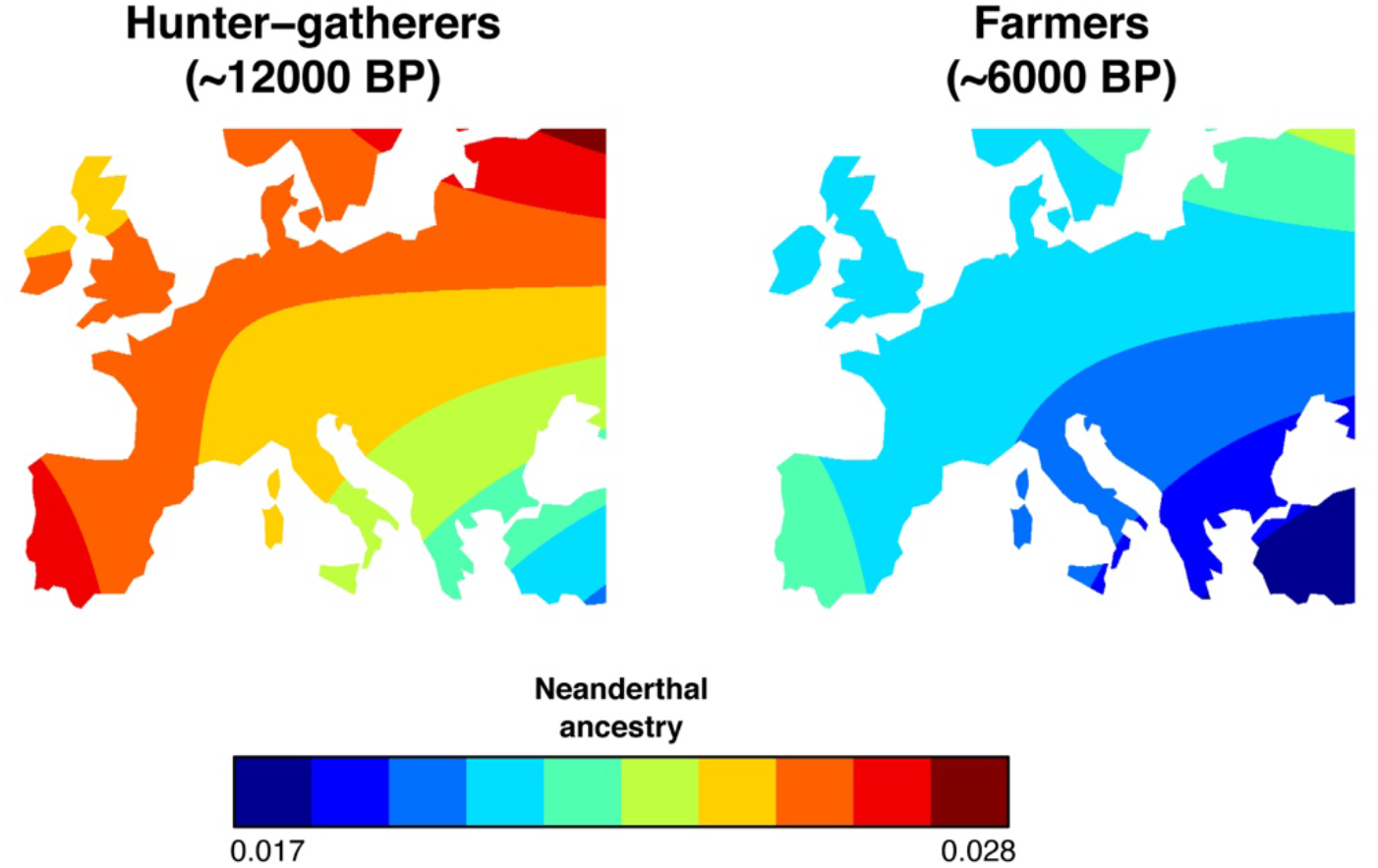
Spatial variation on the level of Neanderthal ancestry in European hunter-gatherers (HG) and farmers (FA) projected using the best “Europe” model (*n* = 1517).

In Europe, both the early human expansion during the Palaeolithic and the farming expansion during the Neolithic trace back to Southwest Asia^30^, where we estimated the lowest level of NE ancestry. The difference in NE ancestry between HGs in Europe and in Anatolia, where FAs originated (see red area and blue area for HGs, respectively, Fig. 5), explains why early FAs contributed to an overall reduction in the level of NE ancestry when expanding across Europe (Fig. 5). Note that it was recently found that modern Levantine and southern Arabian populations still have lower NE ancestry than northern Eurasian populations^49^.

According to the expansion hypothesis^9^, as HGs spread over Europe, their NE introgression levels increased due to a combination of stochastic demographic and migratory processes related to gene surfing^22,23^ and gene dilution^20,30,38^, which explains the spatial gradients of NE ancestry observed in HGs (Fig. 5). Later, a second expansion occurred during the Neolithic, following approximately the same direction as the expansion of HGs. Consequently, the spatial gradient already present in HGs was maintained in FAs because both populations admixed (Fig. 5). As previously noted^44^, the sole observation of a South East to North West genetic cline in Europe is not informative about the expansion of FAs, as the cline could have been generated during the previous HG expansion. Here, our results suggest that the NE ancestry cline was produced during the HG range expansion and was affected by the later expansion of FAs during the Neolithic transition while maintaining the same general orientation. As FAs had initially less NE ancestry than HGs, it lowered the average amount in the admixed European population. This corresponds to the model of population expansion with partial replacement supported by palaeogenomic studies for the Neolithic^31–33^. In the absence of admixture between HGs and FAs, we would have expected a larger decrease in NE ancestry in FA populations, whereas with a large admixture, the gradients in FAs would have rapidly reached that of HGs in Europe.

### Conclusion

Our findings highlight the evolutionary impact of past range expansions in generating spatial gradients of NE ancestry in MHs. We observed that these ancestry levels were more spatially heterogeneous in the past and that they became more homogeneous during the Holocene under the effect of gene flow resulting from population movements and migrations. Complex population movements and genetic interactions are reflected when analysing the level of NE ancestry in different regions (Europe and Asia) and in populations with different cultural backgrounds (hunter-gatherers and farmers). The spatial gradient of NE ancestry in HGs is compatible with a model of range expansion of MHs during the OOA expansion. After this first expansion, the level of NE ancestry was slightly higher in western Eurasia than in eastern Eurasia. Then, a second range expansion of early FAs with a lower NE ancestry than HGs occurred in Europe during the Neolithic transition, from the southeast towards the northwest. This second range expansion is essential for explaining the pattern currently observed of lower NE ancestry in Western Europe than in East Asia^3–6^. Our results therefore do not support our previous hypothesis that the slightly greater NE ancestry in Eastern Eurasia compared to Western Eurasia is due to population dynamics during the OOA expansion of MHs taking place ~ 40000 BP. Instead, our results reveal that the current geographical heterogeneity of NE ancestry is due to dynamics that occurred during the more recent Neolithic expansion ~10,000 BP. The early FA populations, which are probably related to the previously identified basal Eurasian lineage^18,19,49^, admixed with HG, leading to a progressive increase in HG ancestry and consequently NE ancestry in the expanding FA populations. The partial replacement of HGs by FAs^31,32^ thus contributed to reducing the level of NE ancestry in western Eurasia more than in eastern Eurasia. While selection was invoked to explain the difference between Europe and Asia^43^, the neutral hypothesis of historical range expansions is sufficient to explain past and current patterns of NE ancestry in humans.

Although the introgression of archaic material in MHs was probably counterselected during an initial stage^41^, the fact that the amount of NE introgression is relatively stable through time, approximately 2%, suggests that this remaining small portion of introgressed DNA can be considered to evolve generally neutrally. This assumption is also supported by the observation that archaic introgression tends to be rarer in gene-rich regions^7^. However, there are exceptions to this general pattern, with examples of adaptive introgression related to the immune system^50,51^, skin pigmentation^15^ and altitude^52^, providing a better adaptation to local environmental conditions and pathogens^53^. Moreover, in present-day populations, some loci introgressed from archaic hominins appear to influence disease risk, such as neurological, psychiatric, immunological, dermatological and dietary disorders^5,54^, either positively or negatively. It is thus crucial to describe neutral patterns of introgression resulting from human demography and migration to allow better detection of loci under selection (positive or negative) as outliers of the neutral background. It could help to reconstruct the impact of past epidemics on the evolution of human immunity. Additional palaeogenomic data for the most ancient periods combined with alternative modelling methods should allow a more detailed understanding of evolutionary processes leading to similar diversity patterns. These developments would provide a better understanding of both the population dynamics within our species and its interactions with other extinct archaic species, such as Neanderthals and Denisovans.

## Material and Methods

### Dataset

Palaeogenomic data available for Eurasian populations were retrieved from the AADR database (Allen Ancient DNA Resource v50.0^39^). This database includes 10391 genomes. For each genome, we retained the mean date in years before present (BP) reported in the database, which corresponds either to the mean of the 95.4% CI calibrated radiocarbon age or to the mean of the archaeological context range. We only included genomes from individuals located in Eurasia in our analysis, with longitude 34° as a delimiter, i.e., west of 34° is considered Europe in our analyses, while east of 34° is considered Asia. For genomes that have multiple versions in the database, we retained for analysis only the version with the most SNPs. We filtered out putatively contaminated palaeogenomes by using the contamination warning estimated through linkage disequilibrium^55^ (values of *“contamLD_warning”* being either *“Model_Misspecified”* or *“Very_High_Contamination”* were removed) as well as by excluding genomes that contained “contam” or “possible.contam” in their GroupID name. Moreover, we excluded from the downstream analysis all the genomes missing the geographical coordinates of the location of their origin or information about the population group to which they belong.

We assigned each genome to a population group based either on the information provided by the GroupID parameter or on the information provided in the publications that produced these genomes: Palaeolithic and Mesolithic hunter-gatherers (HG, *n*=135), Neolithic and Chalcolithic farmers (FAs, *n*=810), other ancient genomes not belonging to HG and FA (OT, *n*=2327), and modern genomes (MDs, *n*=1192), which are referenced by a date zero in the database. The table S4 in supporting information lists all the genomes used in our analysis.

### Estimation of Neanderthal ancestry

We used F4-ratios as introduced by Reich et al.^56^ to estimate NE ancestry (α) in each genome of interest. It is calculated as the ratio of two F4-statistics generated from five populations, where one population results from the admixture between two others. We followed Petr et al. ^55^ to avoid a bias in the temporal distribution of the F4-statistics. This procedure considers the reflux of NE introgression from European populations into northern and western African populations using the Dinka population from eastern Africa as a sister population (“C”) of the tested genome (“X”) instead of the Yoruba from western Africa. The Altai Neanderthal (“A”) was used as a sister population of the Vindija Neanderthal (“B”), and chimpanzee was included as an outgroup (“O”). The value of α estimates the proportion of ancestry deriving from B in an admixed genome X, using the same A, C and O populations as references: α = F4(A, O, X, C)/F4(A, O, B, C), where X = test genome; A = Altai_Neanderthal. DG; B = Vindija_Neanderthal. DG; C = Dinka. DG; O = Chimp. REF; all retrieved from the AADR database. We refer to this observed F4 ratio as Neanderthal ancestry (or introgression).

We used the R package ADMIXTOOLS 2 to compute the F4-ratio for each genome^57^. Only genomes showing a significant value (Z > 3) were included in our analyses to ensure the accuracy of the estimated NE ancestry. This resulted in 4464 genomes. The European area is represented by 2146 palaeogenomes, and the Asian area is represented by 2318 palaeogenomes. We averaged the F4-ratio for genomes with the same date, geographic coordinates and population group, resulting in a final dataset of 2625 population samples (from ~40000 years BP to modern time, 1517 in Europe and 1108 in Asia, Fig. 1B).

### Statistical analysis

We used the computed F4-ratio as the response variable in a series of linear mixed model (LMM) analyses, which are especially well suited to describe the relation between variables, including autocorrelation and missing data. This response variable was log-transformed to maintain the Gaussian distribution of residuals. In the first analysis, we included latitude, longitude, time (years BP), continental area (Europe or Asia), and their interactions as explanatory (fixed) variables. We evaluated the effect of the population groups (HG, FA, OT and MD), as well as the nested effect of time period within these population groups (i.e., grouping dates within 500 years ranges), as random variables. We also evaluated the influence of the spatial autocorrelation and temporal autocorrelation of the dataset. The choice of fixed and random model structures from among those investigated and the selection of the best model was based on the lowest Akaike Information Criterion value (AIC)^40^, following Zuur et al.^67^. The final model structure was an LMM that considered the period nested in the population groups as a random intercept and slope of continental area, as well as an exponential spatial autocorrelation. We called this model “Full Eurasia” because it incorporates the whole dataset (Table S1, supporting information).

Because the ancient DNA samples belonging to different population groups were not equally distributed throughout Europe and Asia, we ran two additional LMM analyses using subsets of the data to explore in more detail the fixed effect of population groups on the log-transformed F4-ratio. The autocorrelation structure, as well as the random and fixed effects of both models, were selected in a similar way as the “Full Eurasia” model. First, we focused on the temporal trend in Europe and Asia, considering time, continental area, and population groups (HG-FA-OT) as fixed variables, also including their respective interactions in the model. We excluded MD samples because there is no variation allowing a cross-level interaction with time (all dates are identical). The final model structure considered the period (i.e., in clusters of 500 years) as a random intercept and slope of continental area, as well as an exponential spatial autocorrelation. This model is reported as “Ancient Eurasia” because it considers only the ancient genomes from Eurasia as a whole (Table S1, supporting information). Second, because the spatial density of available palaeogenomes (i.e., the number of palaeogenomes per millions of square kilometres) is much larger in Europe (191.55) than in Asia (28.15), we focused the spatial analysis of population groups on Europe. The fixed variables were latitude, longitude, population groups (HG-FA-OT-MD) and their interactions, also including time as a fixed covariate. The interaction between time and population groups was excluded because of the lack of temporal variation in MD samples. The final LMM considered the period as a random intercept, as well as a ratio quadratic spatial autocorrelation. This model is reported as “Europe” (Table S1, supporting information).

For all three LMM analyses, we verified collinearity among fixed variables by computing the variance inflator factor (VIF). All variables included in the separate models did not show collinearity with other variables (VIFs < 4)^58^. We reported the conditional (*R^2^*_GLMM(c)_) and marginal (*R^2^*_GLMM(m)_) coefficients of determination for the selected LMMs (Table S5, supporting information). They denote the proportion of variance explained by fixed variables and both fixed and random variables, respectively^59^. Because the number of samples within continental locations and groups of populations were not the same, we reported mean differences by considering an ANCOVA with sum of squares of type III. A Tukey correction was implemented for multiple post hoc comparisons among population groups. We reported trends of continuous variables by considering the average values of other predictors within continental areas and population groups (Table S6, supporting information), except when it is specifically mentioned in the main text. We show the outputs of the best models after the selection of the random and fixed structure (Table S1, supporting information), but we also show the average of candidate models based on a cumulative weighted AIC of 90% (Σ_*w_i_*_ ≥ 0.90, Table S2, supporting information), as well as the full model without selection of fixed variables (Table S3, supporting information). The last two models were referred to when a nonsignificant relationship was excluded from the best models. All analyses were carried out using the R language^70^. The *nlme* package^71^ was used for the LMMs, the *emmeans* package^73^ for multiple comparisons, and the *MuMIn* package^72^ for the estimation of pseudodetermination coefficients and model averaging.

## Author contribution

All authors contributed to the design of the study, the interpretation of the results and to the writing of the manuscript. MC coordinated the study. JR and AT formatted the data. JR, CQ and AT performed the analyses. JR and CQ drafted the manuscript.

## Acknowledgments

This project was financially supported by the Swiss National Research Foundation grant n° 31003A_182577 to MC and n° P5R5PB_203169 to CSQ. The authors would like to thank Pascale Gerbault and Paola Cerrito and Lionel Di Santo for a careful reading of the manuscript.

## Data availability statement

All the data analysed were previously published and retrieved from the Allen Ancient DNA Resource v50.0 (https://reich.hms.harvard.edu/allen-ancient-dna-resource-aadr-downloadable-genotypes-present-day-and-ancient-dna-data). The table S4 in supporting information lists all the genomes used in our analysis with their indexes in the AADR database and their original references, as well as the associated population groups.

## Code availability

All the R packages used to perform the analyses are listed in the Material and Method section and freely available from the Comprehensive R Archive Network (https://cran.r-project.org/).

## References

1 Green, R. E. et al. A draft sequence of the Neandertal genome. Science 328, 710–722 (2010). https://doi.org:10.1126/science.1188021

2 Reich, D. et al. Genetic history of an archaic hominin group from Denisova Cave in Siberia. Nature 468, 1053–1060 (2010). https://doi.org:10.1038/Nature09710

3 Chen, L., Wolf, A. B., Fu, W., Li, L. & Akey, J. M. Identifying and Interpreting Apparent Neanderthal Ancestry in African Individuals. Cell 180, 677–687 e616 (2020). https://doi.org:10.1016/j.cell.2020.01.012

4 Meyer, M. et al. A high-coverage genome sequence from an archaic Denisovan individual. Science 338, 222–226 (2012). https://doi.org:10.1126/science.1224344

5 Prufer, K. et al. A high-coverage Neandertal genome from Vindija Cave in Croatia. Science 358, 655–658 (2017).

6 Wall, J. D. et al. Higher Levels of Neanderthal Ancestry in East Asians than in Europeans. Genetics 194, 199–+ (2013). https://doi.org:10.1534/genetics.112.148213

7 Sankararaman, S. et al. The genomic landscape of Neanderthal ancestry in present-day humans. Nature (2014). https://doi.org:10.1038/nature12961

8 Villanea, F. A. & Schraiber, J. G. Multiple episodes of interbreeding between Neanderthal and modern humans. Nat Ecol Evol 3, 39–44 (2019). https://doi.org:10.1038/s41559-018-0735-8

9 Quilodrán, C. S., Tsoupas, A. & Currat, M. The Spatial Signature of Introgression After a Biological Invasion With Hybridization. Front Ecol Evol 8 (2020). https://doi.org:10.3389/fevo.2020.569620

10 Currat, M. & Excoffier, L. Strong reproductive isolation between humans and Neanderthals inferred from observed patterns of introgression. Proc Natl Acad Sci U S A 108, 15129–15134 (2011). https://doi.org:10.1073/pnas.1107450108

11 Eriksson, A. & Manica, A. Effect of ancient population structure on the degree of polymorphism shared between modern human populations and ancient hominins. Proc Natl Acad Sci U S A 109, 13956–13960 (2012). https://doi.org:10.1073/pnas.1200567109

12 Fu, Q. M. et al. The genetic history of Ice Age Europe. Nature 534, 200–+ (2016). https://doi.org:10.1038/nature17993

13 Racimo, F., Sankararaman, S., Nielsen, R. & Huerta-Sanchez, E. Evidence for archaic adaptive introgression in humans. Nat Rev Genet 16, 359–371 (2015). https://doi.org:10.1038/nrg3936

14 Kuhlwilm, M. et al. Ancient gene flow from early modern humans into Eastern Neanderthals. Nature 530, 429–+ (2016). https://doi.org:10.1038/nature16544

15 Vernot, B. & Akey, J. M. Resurrecting Surviving Neandertal Lineages from Modern Human Genomes. Science 343, 1017–1021 (2014). https://doi.org:10.1126/science.1245938

16 Prufer, K. et al. The complete genome sequence of a Neanderthal from the Altai Mountains. Nature 505, 43–49 (2014). https://doi.org:10.1038/nature12886

17 Krause, J. et al. Neanderthals in central Asia and Siberia. Nature 449, 902–904 (2007). https://doi.org:10.1038/nature06193

18 Lazaridis, I. et al. Genomic insights into the origin of farming in the ancient Near East. Nature 536, 419–+ (2016). https://doi.org:10.1038/nature19310

19 Vernot, B. & Akey, J. M. Complex history of admixture between modern humans and Neandertals. American journal of human genetics 96, 448–453 (2015). https://doi.org:10.1016/j.ajhg.2015.01.006

20 Barbujani, G., Sokal, R. R. & Oden, N. L. Indo-European origins: a computer-simulation test of five hypotheses. Am J Phys Anthropol 96, 109–132 (1995).

21 Rendine, S., Piazza, A. & Cavalli-Sforza, L. Simulation and separation by principal components of multiple demic expansions in Europe. Am. Nat. 128, 681–706 (1986).

22 Edmonds, C. A., Lillie, A. S. & Cavalli-Sforza, L. L. Mutations arising in the wave front of an expanding population. Proc Natl Acad Sci U S A 101, 975–979 (2004). https://doi.org:10.1073/pnas.0308064100

23 Klopfstein, S., Currat, M. & Excoffier, L. The fate of mutations surfing on the wave of a range expansion. Mol Biol Evol 23, 482–490 (2006). https://doi.org:10.1093/molbev/msj057

24 Peischl, S., Dupanloup, I., Bosshard, L. & Excoffier, L. Genetic surfing in human populations: from genes to genomes. Curr Opin Genet Dev 41, 53–61 (2016). https://doi.org:10.1016/j.gde.2016.08.003

25 Travis, J. M. et al. Deleterious mutations can surf to high densities on the wave front of an expanding population. Mol Biol Evol 24, 2334–2343 (2007). https://doi.org:10.1093/molbev/msm167

26 Austerlitz, F., JungMuller, B., Godelle, B. & Gouyon, P. H. Evolution of coalescence times, genetic diversity and structure during colonization. Theor Popul Biol 51, 148–164 (1997). https://doi.org:10.1006/tpbi.1997.1302

27 Li, J. Z. et al. Worldwide human relationships inferred from genome-wide patterns of variation. Science 319, 1100–1104 (2008). https://doi.org:10.1126/science.1153717

28 Sousa, V., Peischl, S. & Excoffier, L. Impact of range expansions on current human genomic diversity. Curr Opin Genet Dev 29, 22–30 (2014). https://doi.org:10.1016/j.gde.2014.07.007

29 Currat, M., Ruedi, M., Petit, R. J. & Excoffier, L. The hidden side of invasions: Massive introgression by local genes. Evolution 62, 1908–1920 (2008). https://doi.org:10.1111/j.1558-5646.2008.00413.x

30 Currat, M. & Excoffier, L. The effect of the Neolithic expansion on European molecular diversity. Proc Biol Sci 272, 679–688 (2005). https://doi.org:10.1098/rspb.2004.2999

31 Brandt, G. et al. Ancient DNA Reveals Key Stages in the Formation of Central European Mitochondrial Genetic Diversity. Science 342, 257–261 (2013). https://doi.org:10.1126/science.1241844

32 Lipson, M. et al. Parallel palaeogenomic transects reveal complex genetic history of early European farmers. Nature 551, 368–+ (2017). https://doi.org:10.1038/nature24476

33 Silva, N. M., Rio, J., Kreutzer, S., Papageorgopoulou, C. & Currat, M. Bayesian estimation of partial population continuity using ancient DNA and spatially explicit simulations. Evol Appl 11, 1642–1655 (2018). https://doi.org:10.1111/eva.12655

34 Hofmanova, Z. et al. Early farmers from across Europe directly descended from Neolithic Aegeans. Proc Natl Acad Sci U S A 113, 6886–6891 (2016). https://doi.org:10.1073/pnas.1523951113

35 Allentoft, M. E. et al. Population genomics of Bronze Age Eurasia. Nature 522, 167–172 (2015). https://doi.org:10.1038/nature14507

36 Haak, W. et al. Massive migration from the steppe was a source for Indo-European languages in Europe. Nature 522, 207–211 (2015). https://doi.org:10.1038/nature14317

37 Rio, J., Quilodran, C. S. & Currat, M. Spatially explicit paleogenomic simulations support cohabitation with limited admixture between Bronze Age Central European populations. Commun Biol 4, 1163 (2021). https://doi.org:10.1038/s42003-021-02670-5

38 Chikhi, L., Nichols, R. A., Barbujani, G. & Beaumont, M. A. Y genetic data support the Neolithic demic diffusion model. Proc Natl Acad Sci U S A 99, 11008–11013 (2002). https://doi.org:10.1073/pnas.162158799

39 Allen Ancient DNA Resource Version 50. (2022).

40 Burnham, K. P. & Anderson, D. R. A Practical Information-Theoretic Approach. Model Selection and Multimodel Inference. 2nd edn, (Springer, 2002).

41 Petr, M., Paabo, S., Kelso, J. & Vernot, B. Limits of long-term selection against Neandertal introgression. P Natl Acad Sci USA 116, 1639–1644 (2019). https://doi.org:10.1073/pnas.1814338116

42 Nielsen, R. et al. Tracing the peopling of the world through genomics. Nature 541, 302–310 (2017). https://doi.org:10.1038/nature21347

43 Juric, I., Aeschbacher, S. & Coop, G. The Strength of Selection against Neanderthal Introgression. PLoS Genet 12, e1006340 (2016). https://doi.org:10.1371/journal.pgen.1006340

44 Arenas, M., Francois, O., Currat, M., Ray, N. & Excoffier, L. Influence of admixture and paleolithic range contractions on current European diversity gradients. Mol Biol Evol 30, 57–61 (2013). https://doi.org:10.1093/molbev/mss203

45 Arenas, M., Ray, N., Currat, M. & Excoffier, L. Consequences of Range Contractions and Range Shifts on Molecular Diversity. Mol Biol Evol 29, 207–218 (2012). https://doi.org:10.1093/molbev/msr187

46 Fuller, D. Q., Willcox, G. & Allaby, R. G. Cultivation and domestication had multiple origins: arguments against the core area hypothesis for the origins of agriculture in the Near East. World Archaeol 43, 628–652 (2011). https://doi.org:10.1080/00438243.2011.624747

47 Bettinger, R. L., Barton, L. & Morgan, C. The Origins of Food Production in North China: A Different Kind of Agricultural Revolution. Evol Anthropol 19, 9–21 (2010). https://doi.org:10.1002/evan.20236

48 Riehl, S., Zeidi, M. & Conard, N. J. Emergence of Agriculture in the Foothills of the Zagros Mountains of Iran. Science 341, 65–67 (2013). https://doi.org:10.1126/science.1236743

49 Vyas, D. N. & Mulligan, C. J. Analyses of Neanderthal introgression suggest that Levantine and southern Arabian populations have a shared population history. American Journal of Physical Anthropology 169, 227–239 (2019). https://doi.org:10.1002/ajpa.23818

50 Mendez, F. L., Watkins, J. C. & Hammer, M. F. A haplotype at STAT2 Introgressed from neanderthals and serves as a candidate of positive selection in Papua New Guinea. American journal of human genetics 91, 265–274 (2012). https://doi.org:10.1016/j.ajhg.2012.06.015

51 Quach, H. et al. Genetic Adaptation and Neandertal Admixture Shaped the Immune System of Human Populations. Cell 167, 643–+ (2016). https://doi.org:10.1016/j.cell.2016.09.024

52 Huerta-Sanchez, E. et al. Altitude adaptation in Tibetans caused by introgression of Denisovan-like DNA. Nature 512, 194–+ (2014). https://doi.org:10.1038/nature13408

53 Kerner, G., Patin, E. & Quintana-Murci, L. New insights into human immunity from ancient genomics. Curr Opin Immunol 72, 148–157 (2021). https://doi.org:10.1016/j.coi.2021.04.006

54 Zeberg, H. & Paabo, S. The major genetic risk factor for severe COVID-19 is inherited from Neanderthals. Nature 587, 610–+ (2020). https://doi.org:10.1038/s41586-020-2818-3

55 Nakatsuka, N. et al. ContamLD: estimation of ancient nuclear DNA contamination using breakdown of linkage disequilibrium. Genome Biol 21, 199 (2020). https://doi.org:10.1186/s13059-020-02111-2

56 Reich, D., Thangaraj, K., Patterson, N., Price, A. L. & Singh, L. Reconstructing Indian population history. Nature 461, 489–U450 (2009). https://doi.org:10.1038/nature08365

57 Maier, R., Flegontov, P., Flegontova, O., Changmai, P. & Reich, D. Admixtools 2. bioRxiv (2022). https://github.com/uqrmaie1/admixtools>.

58 Zuur, A., Ieno, E. N., Walker, N., Saveliev, A. A. & Smith, G. M. Mixed Effects Models and Extensions in Ecology With R. (Springer Science & Business Media, 2009).

59 Nakagawa, S., Johnson, P. C. D. & Schielzeth, H. The coefficient of determination R-2 and intra-class correlation coefficient from generalized linear mixed-effects models revisited and expanded. J R Soc Interface 14 (2017). https://doi.org:10.1098/rsif.2017.0213

