## supporting information for "Past human expansions shaped the spatial pattern of Neanderthal ancestry"

[†] contributed equally to this work

* corresponding author

**Table S1.** Best models on the level of Neanderthal ancestry in Eurasia. The models were selected based on the lowest AIC values. They evaluate the effect of spatial (latitude and longitude) and temporal (date in years BP) variation on the level of Neanderthal ancestry. Groups of populations are represented by hunter gatherers (HGs), Neolithic farmers (FAs), other ancient samples (OTs), and modern samples (MDs). They were collected in different locations in Europe and Asia. A) Full spatiotemporal dataset in Eurasia (~40000 years BP – current time). B) Temporal subset of the full dataset focusing on ancient DNA samples in Europe and Asia. C) Spatial subset of the full dataset focusing on Europe. Neanderthal ancestry was log-transformed (see methods).

| Model | Variable | Estimate | SE | *t* | p-value |
|---|---|---|---|---|---|
| A) Full Eurasia | Intercept | -4.569 | 0.085 | -53.67 | <0.001 |
| (*n* = 2625) | Latitude | 0.014 | 0.002 | 7.500 | <0.001 |
| | Longitude | 0.006 | 0.001 | 6.119 | <0.001 |
| | Time | -1.8E-05 | 7.3E-06 | -2.405 | 0.016 |
| | Continent:Europe | 0.526 | 0.087 | 6.057 | <0.001 |
| | Latitude x Longitude | -1.1E-04 | 2.1E-05 | -5.086 | <0.001 |
| | Latitude x Continent:Europe | -0.011 | 0.002 | -5.930 | <0.001 |
| | Latitude x Time | 3.0E-07 | 1.4E-07 | 2.099 | 0.036 |
| | Longitude x Continent:Europe | -0.002 | 0.001 | -2.767 | 0.006 |
| | Time x Continent:Europe | 1.1E-05 | 3.0E-06 | 3.745 | <0.001 |
| B) Ancient Eurasia | Intercept | -3.750 | 0.044 | -86.007 | <0.001 |
| (*n* = 2534) | Time | -2.6E-05 | 6.98E-06 | -3.671 | <0.001 |
| | Population:HG | -0.050 | 0.0545 | -0.921 | 0.357 |
| | Population:OT | -0.071 | 0.046 | -1.537 | 0.124 |
| | Continent:Europe | -0.031 | 0.016 | -1.960 | 0.050 |
| | Time x Population:HG | 2.6E-05 | 7.2E-06 | 3.560 | <0.001 |
| | Time x Population:OT | 1.3E-05 | 8.4E-06 | 1.534 | 0.125 |
| | Time x Continent:Europe | 8.0E-06 | 2.6E-06 | 3.037 | 0.002 |
| C) Europe | Intercept | -3.867 | 0.055 | -69.964 | <0.001 |
| (*n* = 1517) | Latitude | -7.9E-04 | 1.1E-03 | -0.724 | 0.469 |
| | Longitude | -0.02 | 3.4E-03 | -5.731 | <0.001 |
| | Time | 7.0E-06 | 2.6E-06 | 2.631 | 0.009 |
| | Population:HG | 0.122 | 0.027 | 4.565 | <0.001 |

| | | | | |
|---|---|---|---|---|
| Population:OT | 0.023 | 0.016 | 1.500 | 0.134 |
| Population:MD | 0.016 | 0.053 | 0.301 | 0.764 |
| Latitude x Longitude | 3.7E-04 | 6.9E-05 | 5.406 | <0.001 |

**Table S2.** Average models on the level of Neanderthal ancestry in Eurasia. The models were averaged based in a cumulative weighted AIC of 90% ($\Sigma\omega_i \geq 0.90$, see Table S5). They evaluate the effect of spatial (latitude and longitude) and temporal (date in years BP) variation on the level of Neanderthal ancestry. Groups of populations are represented by hunter gatherers (HGs), Neolithic farmers (FAs), other ancient samples (OTs), and modern samples (MDs). They were collected in different locations in Europe and Asia. A) Full spatiotemporal dataset in Eurasia (~40000 years BP – current time). B) Temporal subset of the full dataset focusing on ancient DNA samples in Europe and Asia. C) Spatial subset of the full dataset focusing on Europe. Neanderthal ancestry was log-transformed (see methods).

| Model | Variable | Estimate | SE | *z* | p-value |
|---|---|---|---|---|---|
| A) Full Eurasia | Intercept | -4.613 | 0.081 | 56.623 | <0.001 |
| (*n* = 2625) | Latitude | 0.015 | 0.002 | 8.277 | <0.001 |
| | Longitude | 0.006 | 0.001 | 6.223 | <0.001 |
| | Time | -2.6E-06 | 1.7E-06 | 1.594 | 0.111 |
| | Continent:Europe | 0.518 | 0.086 | 6.045 | <0.001 |
| | Latitude x Longitude | -1.1E-04 | 2.1E-05 | 5.154 | <0.001 |
| | Latitude x Continent:Europe | -0.011 | 0.002 | 5.819 | <0.001 |
| | Latitude x Time | 3.0E-13 | 3.3E-10 | 0.001 | 0.999 |
| | Longitude x Continent:Europe | -0.002 | 5.9E-04 | 2.939 | 0.003 |
| | Time x Continent:Europe | 1.1E-05 | 2.9E-06 | 3.720 | <0.001 |
| | Time x Longitude | -4.7E-22 | 8.5E-15 | 0.000 | 1.000 |
| B) Ancient Eurasia | Intercept | -3.75 | 0.044 | 85.935 | <0.001 |
| (*n* = 2534) | Time | -2.6E-05 | 7.0E-06 | 3.669 | <0.001 |
| | Population:HG | -0.050 | 0.055 | 0.921 | 0.357 |
| | Population:OT | -0.071 | 0.046 | 1.537 | 0.124 |
| | Continent:Europe | -0.031 | 0.016 | 1.954 | 0.051 |
| | Time x Population:HG | 2.6E-05 | 7.2E-06 | 3.558 | <0.001 |
| | Time x Population:OT | 1.3E-05 | 8.4E-06 | 1.533 | 0.125 |
| | Continent:Europe x Population:HG | 1.1E-04 | 0.003 | 0.033 | 0.974 |
| | Continent:Europe x Population:OT | 3.4E-05 | 0.001 | 0.024 | 0.980 |
| | Time x Continent:Europe | 8.0E-06 | 2.6E-06 | 3.032 | 0.002 |
| C) Europe | Intercept | -3.867 | 0.056 | 69.075 | <0.001 |
| (*n* = 1517) | Latitude | -8.0E-04 | 0.001 | 0.725 | 0.469 |
| | Longitude | -0.02 | 3.4E-03 | 5.726 | <0.001 |

| | | | | |
|---|---|---|---|---|
| Time | 6.8E-06 | 2.7E-06 | 2.537 | 0.011 |
| Population:HG | 0.125 | 0.047 | 2.657 | <0.001 |
| Population:OT | 0.023 | 0.020 | 1.138 | 0.255 |
| Population:MD | 0.017 | 0.060 | 0.289 | 0.772 |
| Latitude x Longitude | 3.7E-04 | 6.9E-05 | 5.401 | <0.001 |
| Latitude x Time | -3.4E-14 | 1.9E-10 | 0.000 | 1.000 |
| Latitude x Population:HG | -6.2E-05 | 7.4E-04 | 0084 | 0.933 |
| Latitude x Population:OT | 1.5E-05 | 2.3E-04 | 0.063 | 0.950 |
| Latitude x Population:MD | -3.3E-05 | 6.0E-04 | 0.055 | 0.956 |
| Longitude x Time | -4.0E-14 | 1.3E-10 | 0.000 | 1.000 |
| Longitude x Population:HG | -1.1E-10 | 7.1E-07 | 0.000 | 1.000 |
| Longitude x Population:OT | 7.1E-12 | 2.1E-07 | 0.000 | 1.000 |
| Longitude x Population:MD | 7.0E-11 | 8.5E-07 | 0.000 | 1.000 |

**Table S3.** Full models on the level of Neanderthal ancestry in Eurasia with all tested fixed variable. They evaluate the effect of spatial (latitude and longitude) and temporal (date in years BP) variation on the level of Neanderthal ancestry. Groups of populations are represented by hunter gatherers (HGs), Neolithic farmers (FAs), other ancient samples (OTs), and modern samples (MDs). They were collected in different locations in Europe and Asia. A) Full spatiotemporal dataset in Eurasia (~40000 years BP – current time). B) Temporal subset of the full dataset focusing on ancient DNA samples in Europe and Asia. C) Spatial subset of the full dataset focusing on Europe. Neanderthal ancestry was log-transformed (see methods).

| Dataset | Variable | Estimate | SE | *t* | p-value |
|---|---|---|---|---|---|
| A) Full Eurasia | Intercept | -4.569 | 0.085 | -53.626 | <0.001 |
| (*n* = 2625) | Latitude | 0.014 | 0.002 | 7.378 | <0.001 |
| | Longitude | 0.006 | 0.001 | 6.122 | <0.001 |
| | Time | -1.6E-05 | 8.4E-06 | -1.954 | 0.051 |
| | Continent:Europe | 0.526 | 0.087 | 6.060 | <0.001 |
| | Latitude x Longitude | -1.1E-04 | 2.1E-05 | -4.942 | <0.001 |
| | Latitude x Continent:Europe | -0.011 | 0.002 | -5.823 | <0.001 |
| | Latitude x Time | 2.9E-07 | 1.4E-07 | 2.048 | 0.041 |
| | Longitude x Continent:Europe | -0.002 | 5.9E-04 | -2.778 | 0.006 |
| | Longitude x Time | -1.2E-08 | 4.0E-08 | -0.286 | 0.775 |
| | Time x Continent:Europe | 1.0E-05 | 4.0E-06 | 2.595 | 0.010 |
| B) Ancient Eurasia | Intercept | -3.738 | 0.048 | -77.140 | <0.001 |
| (n = 2534) | Time | -2.6E-05 | 7.2E-06 | -3.612 | <0.001 |
| | Population:HG | -0.096 | 0.068 | -1.428 | 0.153 |
| | Population:OT | -0.084 | 0.050 | -1.669 | 0.095 |
| | Continent:Europe | -0.041 | 0.030 | -1.376 | 0.169 |
| | Time x Population:HG | 2.7E-05 | 7.3E-06 | 3.724 | <0.001 |
| | Time x Population:OT | 1.3E-05 | 8.4E-06 | 1.567 | 0.117 |
| | Continent:Europe x Population:HG | 0.052 | 0.052 | 1.004 | 0.316 |
| | Continent:Europe x Population:OT | 0.016 | 0.025 | 0.620 | 0.536 |
| | Time x Continent:Europe | 7.0E-06 | 3.7E-06 | 1.950 | 0.051 |
| C) Europe | Intercept | -3.575 | 0.177 | -20.228 | <0.001 |
| (*n* = 1517) | Latitude | -0.007 | 0.004 | -1.818 | 0.069 |

| | | | | |
|---|---|---|---|---|
| Longitude | -0.021 | 0.004 | -4.878 | <0.001 |
| Time | -4.2E-05 | 2.9E-05 | -1.447 | 0.148 |
| Population:HG | 0.608 | 0.210 | 2.900 | 0.004 |
| Population:OT | -0.197 | 0.131 | -1.502 | 0.133 |
| Population:MD | -0.033 | 0.306 | -0.108 | 0.914 |
| Latitude x Longitude | 3.8E-04 | 7.5E-05 | 5.070 | <0.001 |
| Latitude x Time | 1.0E-06 | 5.8E-07 | 1.608 | 0.108 |
| Latitude x Population:HG | -0.009 | 0.004 | -2.042 | 0.041 |
| Latitude x Population:OT | 0.004 | 0.003 | 1.637 | 0.102 |
| Latitude x Population:MD | -4.9E-04 | 0.006 | -0.077 | 0.939 |
| Longitude x Time | 1.9E-07 | 3.0E-07 | 0.638 | 0.524 |
| Longitude x Population:HG | -0.004 | 0.002 | -1.570 | 0.117 |
| Longitude x Population:OT | 5.2E-04 | 0.001 | 0.379 | 0.704 |
| Longitude x Population:MD | 0.005 | 0.005 | 1.045 | 0.296 |

**Table S4.** Genomes included in the present study. Population group: group associated to each sample depending on its cultural background and age (HG, FA, OT, MD). Continental region: if the sample was located in Asia or Europe. Master IDs: the Master ID in the AADR database of all the genomes of the sample. Indices: The indices in the information files of the AADR database of all the genomes included in the sample. Publications: the publications of the studies that produced the genomes (see annex file).

**Table S5.** Candidate models to explain the level of Neanderthal ancestry in Eurasia. The model selection considered a cumulative weighted AIC of 90% ($\Sigma\omega_i \geq 0.90$).

| Model title | Models with their variables* | df | AIC | ΔAIC | $\omega_{AIC}$ | $R^2_{GLMM(m)}$ | $R^2_{GLMM(c)}$ |
|---|---|---|---|---|---|---|---|
| Full Eurasia | time+lat+continent+long+time:lat+time:continent+lat:continent+lat:long+continent:long | 19 | -1612.361 | 0.000 | 0.484 | 0.082 | 0.147 |
| | time+lat+continent+long+time:lat+time:continent+time:long+lat:continent+lat:long+continent:long | 20 | -1610.441 | 1.920 | 0.185 | 0.082 | 0.147 |
| | time+lat+continent+long+time:continent+lat:continent+lat:long+continent:long | 18 | -1610.277 | 2.085 | 0.171 | 0.079 | 0.146 |
| Ancient Eurasia | time+continent+pop+time:continent+time:pop | 14 | -1380.770 | 0.000 | 0.663 | 0.035 | 0.043 |
| | time+continent+pop+time:continent+time:pop+continent:pop | 16 | -1378.090 | 2.681 | 0.173 | 0.036 | 0.044 |
| Europe | time+lat+long+pop+lat:long | 12 | -955.368 | 0.000 | 0.162 | 0.072 | 0.097 |
| | time+lat+long+pop+time:lat+lat:long+lat:pop | 16 | -955.236 | 0.132 | 0.151 | 0.077 | 0.102 |
| | time+lat+long+pop+lat:long+lat:pop | 15 | -954.884 | 0.484 | 0.127 | 0.076 | 0.098 |
| | time+lat+long+pop+time:long+lat:long | 13 | -953.980 | 1.388 | 0.081 | 0.072 | 0.097 |
| | time+lat+long+pop+time:lat+time:long+lat:long+lat:pop | 17 | -953.571 | 1.797 | 0.066 | 0.077 | 0.102 |
| | time+lat+long+pop+time:lat+lat:long | 13 | -953.509 | 1.859 | 0.064 | 0.072 | 0.096 |
| | time+lat+long+pop+time:long+lat:long+lat:pop | 16 | -953.420 | 1.948 | 0.061 | 0.076 | 0.098 |
| | time+lat+long+pop+lat:long+long:pop | 15 | -952.670 | 2.698 | 0.042 | 0.073 | 0.095 |
| | time+lat+long+pop+time:lat+lat:long+lat:pop+long:pop | 19 | -952.524 | 2.844 | 0.039 | 0.078 | 0.100 |
| | time+lat+long+pop+lat:long+lat:pop+long:pop | 18 | -952.327 | 3.041 | 0.035 | 0.077 | 0.097 |
| | time+lat+long+pop+time:lat+time:long+lat:long | 14 | -952.146 | 3.221 | 0.032 | 0.072 | 0.098 |
| | lat+long+pop+lat:long+lat:pop | 14 | -951.687 | 3.681 | 0.026 | 0.074 | 0.091 |

*Code of each variable: Latitude (lat), Longitude (long), Sample age in YBP (time), Population group (pop), Continental area (continent).

**Table S6.** Fixed variables considered in the analysis of spatiotemporal variation in Neanderthal ancestry. Average values and standard deviations (in brackets) are presented. HGs: hunter gatherers, FAs: Neolithic farmers, OTs: other ancient samples, MDs: modern samples. Date is presented in years before present (BP).

| | Europe | | | | | Asia | | | | |
|---|---|---|---|---|---|---|---|---|---|---|
| | HG | FA | OT | MD | All | HG | FA | OT | MD | All |
| Latitude | 48.59 (5.65) | 47.73 (5.58) | 49.47 (6.43) | 49.16 (9.17) | 48.83 (6.19) | 49.97 (9.07) | 42.56 (7.55) | 44.97 (7.95) | 34.19 (13.05) | 44.02 (8.84) |
| Longitude | 17.28 (9.66) | 9.18 (11.3) | 11.46 (8.43) | 14.77 (10.83) | 11.11 (9.77) | 83.95 (43.53) | 84.16 (35.68) | 75.39 (24.04) | 84.69 (31.27) | 77.65 (27.66) |
| Date | 10779 (6310) | 5911 (1080) | 3132 (1342) | 0 (0) | 4504 (2906) | 15752 (11888) | 6326 (1544) | 2935 (1309) | 0 (0) | 3657 (3536) |

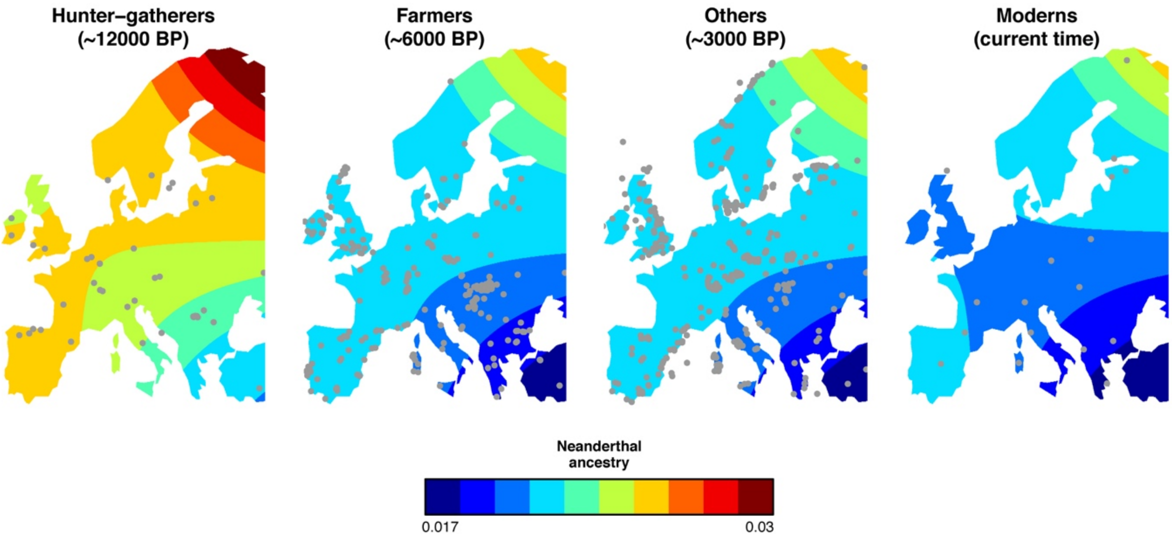

**Figure S1**. Spatial variation on the level of Neanderthal ancestry in different population groups across time expected using the best “Europe” model (n = 1517). The grey dots represent the distribution of DNA samples.
